# Single-Cell Proteomics Reveals Heterogeneous and Bimodal Proteome Responses to DNA Damage

**DOI:** 10.64898/2026.07.20.739584

**Authors:** Kish R. Adoni, Georgina H. Charlton, Jonathan E. Ditcham, Declan T. Cook, Jenny Ho, Joanna Kirkpatrick, Riccardo Zenezini Chiozzi, Konstantinos Thalassinos

## Abstract

DNA damage response (DDR) involves coordinated activation of repair, checkpoint, and stress-adaptive signalling pathways following genomic toxicity. Bulk-cell proteomics, however, averages divergent responses across neighbouring cells, obscuring the characterisation of biologically meaningful proteome remodelling during genotoxic stress. This is particularly relevant to DDR, where a subset of cells undergo apoptosis, resulting in the sampling of dead cells that confound downstream biological interpretation. Here, we applied single-cell proteomics (SCP) to investigate the proteome response of U-2 OS cells to 6-thioguanine (6TG)-induced DNA damage. By filtering individual cells according to predefined morphological-, sampling- and cell-cycle state-criteria, to ensure a matched comparison between control and DNA-damaged populations; 6TG-treatment was found to activate canonical DDR, including mismatch repair and base excision repair, together with secondary stress responses linked to oxidative stress, mitochondrial dysfunction, inflammatory signalling and proteostasis. Underscoring this, DNA-mismatch repair protein MSH3 and mitochondrial energy metabolism regulator PKRAB1, underwent the 4^th^ and 3^rd^ greatest increased protein abundances following DNA-damage, respectively. We then exploited the individual cell resolution of SCP to interrogate neighbouring cell protein-abundance heterogeneity, revealing significant globally increased protein abundance variability following DNA-damage. In particular, proteins involved in mitochondrial oxidative phosphorylation and the tricarboxylic acid cycle exhibited a striking increase in cell-to-cell abundance variability, likely reflecting heterogeneity in the secondary stress responses elicited across U-2 OS cells following 6TG-treatment. Furthermore, analysis of protein-abundance heterogeneity revealed proteins that exhibit bifurcated abundance states across neighbouring cells, with DNA damage inducing either the emergence or abolishment of these bimodal high-low abundance distributions. Indeed, 6TG treatment was associated with remodelling of high–low abundance subpopulation structure across several DNA damage response (PMS1, ORC6), inflammatory (LGALS3, IKKα), metabolic (IDH3A) and proteostasis-related (STAMBP, NAA30, KLHDC4) proteins, indicating that DNA damage can reorganise cell-to-cell protein abundance distributions towards either distinct or more homogeneous proteomic states. Collectively, these findings demonstrate how SCP reveals cell-to-cell variation in the proteomic response to DNA damage that is obscured by population-level averaging.

## INTRODUCTION

DNA damage response (DDR) is a coordinated cellular process that detects genomic lesions and activates repair pathways^1,2^. Maintaining genome integrity requires the integration of multiple repair mechanisms and signalling pathways, including cell-cycle checkpoint signalling, DNA mismatch repair (MMR) and homologous recombination (HR)^2,3^. Disruption of these pathways contributes to genome instability and disease, whilst controlled activation is vital to the maintenance of genome integrity^1^. Understanding the coordinated protein machinery that orchestrates the cellular DDR therefore represents an important challenge towards accurately characterising DDR-associated diseases such as cancer and neurodegeneration.

Thiopurines, such as 6-thioguanine (6TG), represent a class of DNA-damaging compounds that exert cytotoxicity primarily through their incorporation into genomic DNA during replication^4,5^. Following incorporation, 6TG residues are methylated *in-situ* and subsequently recognized by the MMR machinery, triggering iterative repair cycles that generate replication stress and DNA double-strand breaks^4,6^ (Fig. 1A). In response, cells typically undergo ATR– CHK–mediated checkpoint arrest during the S- or G2/M- phase of the cell cycle, followed by HR–mediated repair of replication-associated double-strand breaks, or apoptosis if the damage cannot be resolved^7^. Accordingly, investigation of proteome remodelling following 6TG-treatment provides a tractable system for probing coordinated proteome-level responses to DNA damage^8–10^. Proteomics approaches have played an important role in defining signalling networks activated downstream of DNA damage^10–12^. Most previous studies have relied on bulk-cell measurements that average responses across heterogeneous populations. Because genotoxic stress induces variable responses across individual cells, including differences in repair capacity, checkpoint activation, and commitment to cell death^2,13^; population-level measurements can obscure biologically meaningful subpopulation-specific responses.

**Fig. 1.**
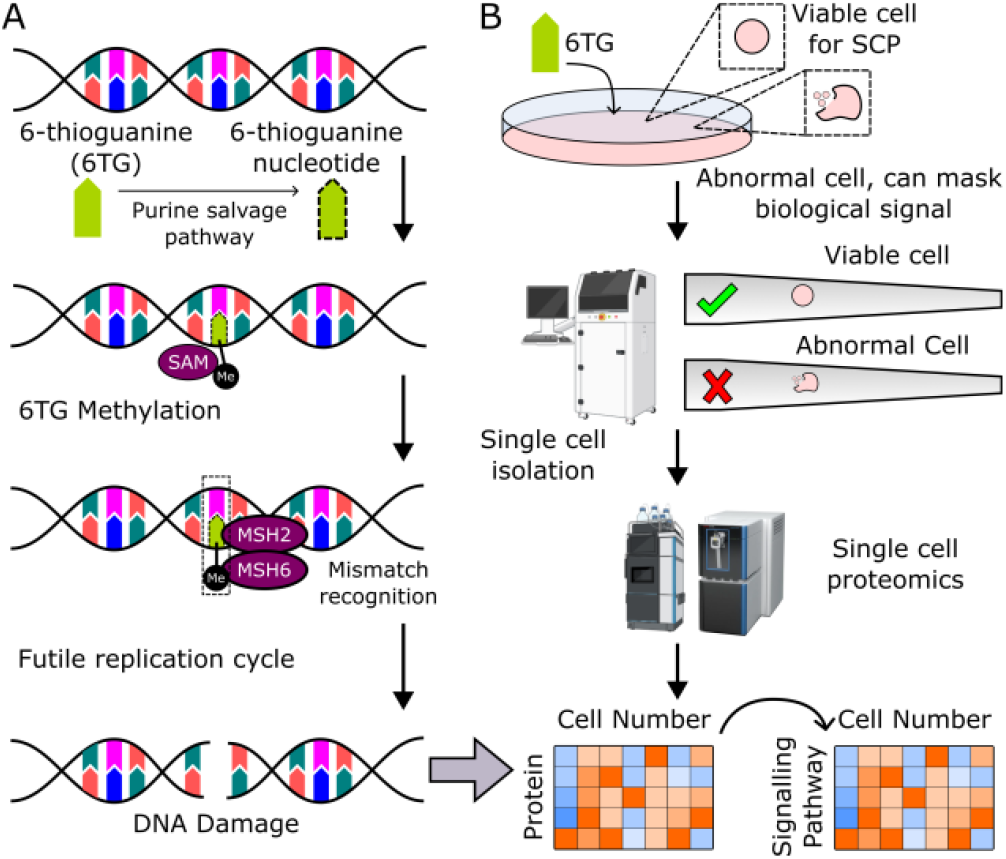
Single-cell proteomics to investigate DNA-damage response. (A) Schematic representation of mechanism of action for 6-thioguanine. After incorporation into genomic DNA and subsequent methylation, 6TG miss-pairs with thymine during the next round of replication. Although the mismatch repair (MMR) machinery recognizes this mismatch and excises the newly synthesized daughter strand, the template strand still contains the modified 6TG base, leading to a ‘futile replication cycle’ of excision and resynthesis that generate persistent DNA lesions. (B) The removal of abnormal cells that don’t satisfy pre-defined morphological thresholds, prior to single cell proteomics analysis, alleviates the impact of such damaged, late-stage apoptotic and/or dead cells from onward analysis, thus preventing their proteome from obscuring the specific DNA-damage response and secondary pathway signatures that are activated following genotoxicity. The onward analysis of these filtered cells thus reveals a clearer picture of their proteome, and consequently, their distinct protein-derived cellular profiles upon DNA-damage.

Recent advances in single-cell proteomics (SCP) now enable direct quantification of proteome remodelling at individual-cell resolution^13,14^. Because DNA damage frequently generates distinct cellular subpopulations; for example, S-phase versus G2/M checkpoint arrest, or DDR engagement versus apoptosis^2^; SCP provides an opportunity to selectively analyse defined cellular states and alleviate signal masking caused by averaging across heterogeneous populations (Fig. 1B). Furthermore, by capturing cell-to-cell variation in protein abundance across thousands of proteins simultaneously, SCP enables unbiased interrogation of stress-response programs with associated changes in the heterogeneity of these responses from cell-to-cell^15^.

Here, we applied SCP to characterize the proteome-wide response of U-2 osteosarcoma (U-2 OS) cells to 6TG-induced DNA damage. This approach, on a focused G1 cellular sub-population undergoing DNA damage response to genotoxic stress, revealed coordinated activation of MMR, nucleotide-excision repair (NER) and base-excision repair (BER) pathways. To this end, DNA mismatch repair protein MSH3 underwent the 4^th^ greatest increased abundance following DNA-damage. In addition to canonical DDR signalling, cells exhibited secondary responses consistent with increased reactive oxygen species (ROS), including activation of oxidative stress defence pathways (NRF2–KEAP1 and non-canonical NF-κB signalling), increased ribosomal stalling, mitochondrial dysfunction, and protein ubiquitination, together representing the cellular trajectory from DNA damage to ROS-activation and proteostatic stress response. By quantifying protein abundance at single-cell resolution, our approach enabled us to examine cell-to-cell heterogeneity in protein abundance across neighbouring cells, revealing a significant increase in the abundance heterogeneity of the cellular proteome following 6TG-induced DNA damage. Interestingly, mitochondrial aerobic respiration related proteins were particularly prone to increased heterogeneity, likely on account of variable levels of mitochondrial dysfunction upon elevated ROS from cell-to-cell.

Our analysis also revealed the bifurcation of protein abundances for some proteins, whereby, rather than exhibiting a continuous distribution of heterogeneity around a mean abundance across neighbouring cells, individual cells segregated into two discrete subpopulations characterized by either high or low relative abundance, with few cells displaying intermediate abundance levels. To this end, several proteins including: STAMBP, NAA30, LGALS3 and PMS1, across the DNA-damage to secondary stress response signalling axis, exhibited the emergence of bifurcation to high-low protein abundances following genotoxic stress. Conversely, some proteins displayed abolishment of intrinsic abundance bimodality (in untreated cells) upon 6TG-treatment, towards a more homogenous distribution of protein abundances. Together, this work demonstrates how single-cell proteomics can uncover heterogeneous proteomic responses to DNA damage that are masked at the population level, enabling a more refined characterisation of the DNA damage response.

## MATERIALS AND METHODS

### Sample Preparation

U-2 OS cells were cultured at 37°C and 5% CO_2_ in Dulbecco’s Modified Eagle’s Medium (DMEM) supplemented with 10% FBS and 1x penicillin-streptomycin. Cells were treated for 16 hours with 6 μM 6TG in PBS; a second batch was kept untreated as a control. After trypsinisation with 0.05% Trypsin-EDTA, cells were washed with phosphate-buffered saline (PBS).

All cells were resuspended in PBS at 150-200 cells/µL for isolation within the CellenONE®. Sample lysis and digestion was performed within a 384-well plate (Thermo Scientific− Surestart) inside the CellenONE. 300 drops of a master mix containing 0.2% DDM, 100mM TEAB and 10ng/µL Trypsin Gold (Promega), was predispensed into the wells. To limit evaporation, humidity was set to 85% and plate temperature was set to 8°C. Individual cells of a diameter between 19 - 22µm and a maximum elongation of 2.3µm were sorted by the CellenONE into the respective wells. This was followed by incubation for 2 h at 50°C with 85% relative humidity inside the instrument. Samples were kept hydrated every 15 min by automated addition of 315 drops of water to each well. After lysis and digestion, samples were cooled to 20°C and 3.2µL of 0.1% TFA was added to the respective wells for quenching and storage. The remaining cells which were not isolated for single-cell analysis were taken for bulk proteomic analysis to create libraries to increase peptide identification. Cells were centrifuged and PBS was removed. Cells were resuspended in 5% SDS and heated to 95°C for 10 minutes to lyse cells. A BCA was performed to establish protein concentration and 100 µg of protein was taken for digestion. A methanol-chloroform precipitation was performed and samples were resuspended in 50 mM ammonium bicarbonate, 1% SDC. Trypsin was added to samples and they were left to digest overnight at 37°C. Reaction was quenched with TFA and SDC was removed. Sample clean up was performed using a Waters Oasis plate C18, samples were dried and resuspended in 0.1% TFA. For libraries, 10 ng of peptide was injected.

### LC-MS/MS

Samples were subjected to the Vanquish Neo UHPLC system (ThermoFisher Scientific) with Buffer A (0.1% formic acid) and Buffer B (acetonitrile, 0.1% formic acid). Peptides were separated on an Aurora Elite column (15×75 XT from IonOpticks) via direct injection. The column was operated at 50°C and connected to an Orbitrap Astral Mass Spectrometer via an EASY-Spray ion source (ThermoFisher Scientific) at a voltage of 1.5kV, with High Field Asymmetric Waveform Ion Mobility Spectrometry (FAIMS) set to a compensation voltage of -50V. Peptides were loaded onto the column at 500nL/min with an elution gradient of 4% to 12% Buffer B over 1.5 minutes, before the flow rate was dropped to 200nL/min over a elution gradient of 12% to 28.5% Buffer B over 8 minutes, then 28.5% to 40% over 1.5 minutes, followed by an increase to 99% at 300nL/min over 1 minute. The Orbitrap Astral MS was operated with an MS1 mass range of 400-800m/z, maximum injection time set to Auto, mass resolution set to 240,000 and an automated gain control (AGC) of 500%. Data Independent Acquisition was applied with an isolation window of 20Th, normalised higher energy collision induced dissociation (HCD) set to 25%. MS2 scans were detected in the Astral mass analyser with an AGC target value of 800% and maximum injection time set to 40ms.

### Data Processing and Analysis

RAW files were processed with Spectronaut v20.5 against the human reference proteome (version 08/20/2023), with trypsin proteolytic enzyme, using the directDIA functionality with standard settings. For this analysis, a “mini-bulk” of 10ng was injected into the LC-MS/MS to acquire a bespoke spectral library to improve protein identification/quantitation. The processed data was then filtered to remove cells that were visualised as abnormal, based on their image from the CellenONE isolator (CellenION). Further, any cell that generated fewer than 2,800 quantified proteins (Fig. 2b) was removed from onward analysis and proteins detected in fewer than 33.3% of cells were also removed. Imputation was performed with PIMMS version 0.5.0^16^. Each experimental condition was imputed independently. Prior to imputation, data was log_2_ transformed and median normalised. PIMMS was run with default Variational Autoencoder model parameters. Cell cycle phase assignment was performed using the marker-based scoring approach implemented in Seurat V5^17,18^. Cell cycle phase was inferred for each cell from the abundance of cell cycle-dependent marker proteins. Markers were taken from the Human Protein Atlas cell cycle-dependent protein list^19^. The S-phase set comprised three proteins (UniProt: P49736, Q03468, Q7L311), and the G2/M set comprised 19 proteins (O14965, O75330, O95067, P46013, P49454, Q02241, Q14008, Q15398, Q8WWK9, Q96GD4, Q99661, Q9BXS6, Q9BZD4, Q9NTJ3, Q9UK76, Q9ULW0, Q9Y6A5, P31350, P52732). Each cell was independently scored for abundance of the S-phase and G2/M-phase sets using the approach of Tirosh et al^20^. For each set, the score was calculated as the average normalised abundance of the marker proteins, minus a background score derived from randomly sampled control proteins of similar abundance. Cells were then assigned to one of three cell cycle phases. Cells with negative scores for both sets were classified as G1, and the remaining cells were assigned to S or G2/M according to the higher score. Only G1-phase cells were used for downstream analysis.

**Fig. 2.**
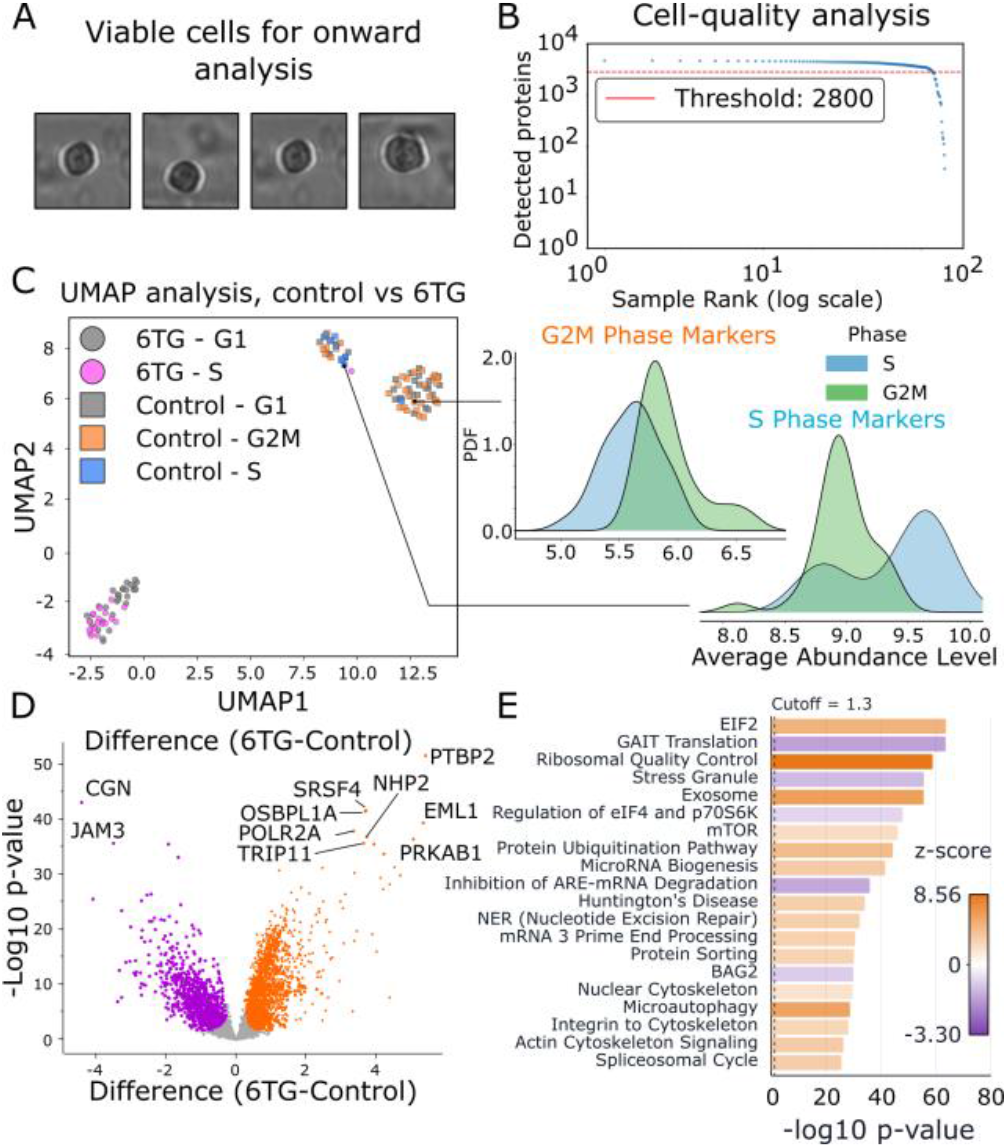
Filtering strategy to improve biological representation. (A) Images acquired from CellenONE single-cell isolator, of individual cells that were dispensed into proteomics sample preparation mixture. Abnormal cells, that did not satisfy pre-selected morphology parameters, were filtered out by CellenONE isolator. (B) Knee-plot to demonstrate the number of quantified protein identifications for each sampled single-cell. A threshold of 2800 quantified proteins was set, such that any cell that did not satisfy this threshold was removed from data processing. (C) UMAP representation of single cell analysis for all processed data, following removal of sub-optimal cells (from morphology and sampling quality). Each cell was then subjected to Seurat cell-cycle analysis, to reveal the distribution of pre-selected markers for S-, G2M- and G1-phase cell-cycle states. From this analysis, each cell was assigned an inferred cell-cycle state, and only those cells that occupied a G1-proteome were retained for further analysis. PDF: Probability Density Function. (D) Volcano plot representation of single-cell proteomics data, for 6TG treated vs untreated cells with statistically significant upregulated proteins upon 6TG-treatment (orange) and statistically significant downregulated proteins upon 6TG-treatment (purple). Top 10 statistically significant proteins are labelled. (E) Top 20 signalling pathways (relative to statistical significance), upon 6TG treatment, with direction of change (z-score) represented by bar colour.

Biological inference was performed using Ingenuity Pathway Analysis (QIAGEN), whereby 3,906 proteins, with their corresponding log_2_ foldchange and adjusted p-value (Benjamini-Hochberg) were analysed. Gene Ontology Cell Component^21^ and WikiPathways^22^ enrichments were acquired using with Cytoscape version 3.10.4^23^. Heterogeneity and bimodality analysis was performed using an in-house developed python script (available at github: https://github.com/kish-adoni/SCP_heterogeneity-and-bimodality or via https://github.com/orgs/ThalassinosLab/repositories).

Individual protein heterogeneity across all cells was calculated from the log_2_ IBAQ using median absolute deviation (MAD):

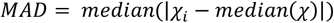

and change in heterogeneity (6TG-control) was calculated via difference in median absolute deviation:

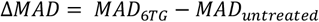

Statistically significant changes in protein abundance heterogeneity following 6TG-treatment were assessed using the Brown-Forsythe test^24^ (median-centred Levene’s test). For each protein, each cell’s abundance was converted to its absolute deviation from the median abundance within its condition, and a one-way analysis of variance was applied to these deviations to test whether their means differed between conditions. P-values were adjusted across tested proteins using the Benjamini– Hochberg correction, with adjusted p-values <0.05 considered significant. Protein abundance multimodality across single cells was assessed independently in untreated and 6-thioguanine (6TG) conditions using Hartigan’s dip test^25^ to quantify deviation from unimodality. For each protein, abundance distributions were additionally modelled using a two-component Gaussian mixture model (GMM) to estimate the proportions of cells occupying each abundance state and the separation between modes. To prioritise biologically meaningful bimodality rather than outlier-driven structure, a balanced bimodality score was computed as:

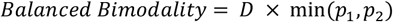

Where *D* is the Hartigan dip statistic and *p*_*1*_, *p*_*2*_ are the fractions of cells assigned to each abundance mode. Proteins were only considered to be bimodal if both abundance modes across the protein satisfied the following criteria; minimum cells per condition: 29, minimum fraction per mode: 0.30, minimum cells per mode: 10, minimum mode separation: 0.5 (log_2_ fold-difference between modes ∼1.4), maximum Dip test p-value: 0.05, minimum valley depth: 0.70, requires 2-component GMM better than 1-component: TRUE. Treatment dependant changes in bimodality were quantified as:

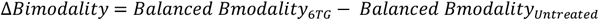

## RESULTS AND DISCUSSION

### Targeted single-cell filtering resolves proteomic responses to DNA-damage

Accurate characterisation of proteome remodelling following DNA damage is complicated by population heterogeneity^1^. Upon induction of genomic lesions, cells activate coordinated DNA-damage response pathways, including MMR and HR, to resolve the damage^2^. Importantly, not all cells successfully complete repair; a subset accumulates unresolved lesions and instead initiates apoptosis and subsequent cell-death^26^, thereby preventing the propagation of damaged DNA to daughter cells. This represents a major challenge to probing the specific proteome response to genomic lesions, which would ordinarily be significantly skewed by the late-stage apoptotic and dead-cell proteome signature of this cellular sub-population. To mitigate this, we applied CellenONE isolation parameters that would identify and discount cells that demonstrated a non-viable morphology, consistent with loss of membrane integrity associated with late-stage apoptosis or cell death (Fig. 2A, S1). Further, such stringency in cell-selection reduced the sampling of doublets that would further skew the data. We next assessed single-cell sampling quality by investigating the number of quantified proteins per cell. An inflection point was observed at approximately 2,800 quantified proteins, below which protein identifications declined sharply. Cells with fewer than 2,800 quantified proteins were therefore excluded from downstream analysis, as reduced proteome coverage is consistent with compromised cellular integrity, cell-death and/or suboptimal mass spectrometric sampling (Fig. 2B).

Beyond cell death via late-stage apoptosis, cell-cycle state is a major source of heterogeneity that can mask biologically relevant proteomic signals, especially in the context of DNA-damage response^27^. Even after synchronisation, proliferative cell cultures can remain enriched across cell-cycle phases^27,28^, and because each phase is associated with distinct proteomic profiles, mixed populations can confound interpretation of condition-specific responses. To mitigate such an effect, we performed a *post-hoc* Seurat cell-cycle analysis against each individual cellular proteome to assign each cell to a cell-cycle state^17^. While this approach infers cell-cycle state from protein abundance rather than directly measuring cell-cycle function, we reasoned that excluding cells assigned to divergent cell-cycle states should reduce cell-cycle composition bias and enable more accurate biological comparisons between conditions. Untreated cells exhibited a heterogeneous distribution of inferred cell-cycle states, comprising G1 (41%), G2/M (46%) and S-phase (9%) populations. Following 16 h of 6TG treatment, this distribution shifted significantly, with all cells assigned to either G1 (62%) or S-phase (38%) and no cells assigned to G2/M. These findings are consistent with 6TG-induced replication stress causing S-phase slowing or arrest^29^. Because 6TG lesions are encountered during DNA replication, MMR-dependent futile repair cycling can activate ATR–CHK1 signalling, suppress replication progression, and prevent efficient transition into G2/M^30^. Thus, the apparent absence of G2/M cells likely reflects accumulation of damaged cells in an S or G1-phase-like proteomic state. Our Seurat-based analysis therefore highlights the importance of accounting for cell-cycle state when comparing across conditions, as differences in the underlying distribution of cell states could otherwise confound biological effects attributed specifically to 6TG-induced DNA damage.

Since 29 cells from the 6TG-treated, and untreated condition, were assigned to the G1-phase, onward analysis was performed by removing all non-G1 cells. To assess the impact of this filtering step; we performed a pathway analysis for pre-vs post-Seurat cell-cycle filtering to reveal the main biological processes that were impacted upon filtering for matched cell cycle states (G1) (Fig. S2). As such, this analysis captured pathway-level changes arising from cell-cycle matching, by comparing the G1-matched 6TG-treated and untreated populations against the original unfiltered comparison, in which untreated cells comprised a heterogenous mixture of inferred cell-cycle states (with G2/M-assignment most common), whereas 6TG-treated cells abolished G2/M-assigned cells. Analysis of the pathways most strongly altered following cell-cycle filtering revealed attenuation of G2/M DNA damage checkpoint regulation, cell-cycle related-FOXO signalling and BTG-mediated cell-cycle regulation, consistent with the removal of cell-cycle-state heterogeneity between conditions (particularly from untreated G2/M cells). Conversely, the increased relative z-scores for Mitotic Prometaphase and Mitotic Roles of Polo-Like Kinase may reflect reduced cell-cycle compositional bias, whereby exclusion of G2/M-phase untreated cells attenuated the 6TG-associated suppression of these pathways^31^. The increased relative contributions of Necrosis^32,33^ and SUMOylation of DNA Replication Proteins may reflect a clearer resolution of genotoxic stress responses specifically associated with 6TG-treatment. These findings highlight the importance of controlling for cell-cycle heterogeneity, thereby removing proteomic signatures associated with differential cell-cycle states that would otherwise mask biologically meaningful proteome-states.

Having filtered the dataset, such that quantified protein abundance changes were more representative of cell-cycle state matched (G1) DDR-associated biological response; we next performed a pathway enrichment analysis of cells, following 6TG-induced DNA-damage (Fig. 2D, E). Canonical DNA damage-associated signalling was upregulated alongside extensive secondary stress-adaptive proteome remodelling. Consistent with the established mechanism of thioguanine-mediated genotoxicity, oxidative stress triggered BER signalling was significantly enriched following 6TG exposure, supporting activation of DNA repair processes downstream of MMR dependent futile repair cycling^34,35^. In parallel, Huntington’s disease– associated signalling was enriched, supporting previous observations that 6TG-induced MMR activation recapitulates proteome features linked to neurodegenerative stress responses^36^. The upregulation of EIF2 signalling^37^ and ribosomal quality control^38^, together with suppression of GAIT translational signalling (suppresses translation of specific inflammatory and stress-response mRNAs^39^), and eIF4/p70S6K signalling (a core protein synthesis and cell growth pathway^40^), indicated extensive reprogramming of the cellular protein synthesis machinery in response to DNA damage–induced stress. Furthermore, enrichment of stress granule signalling, exosome-associated pathways, autophagy, and protein ubiquitination indicated activation of proteostatic stress responses consistent with increased ROS generated following DNA damage.

### 6TG drives secondary ROS-mediated stress response to DNA-damage

Having filtered the single-cell dataset to remove confounding subpopulations that could obscure biologically meaningful proteome shifts, we next investigated the cellular trajectory from primary DNA-damage response through to secondary stress adaptation. By overlaying protein abundance log_2_ fold-changes, following 6TG treatment, onto the canonical MMR pathway (Fig. 3A), our data revealed distinct MMR activation, via increased abundance of MSH2 (log_2_ FC 0.95), MSH3 (4.70), MSH6 (0.67), PCNA (0.61) and LIG1 (0.60), as well as other classical MMR proteins. These findings align with the well characterised role of the MMR machinery in identifying DNA-lesions and initiating DNA-damage response^3^ (Fig. 1). Perhaps surprisingly however, MSH3 was found to undergo the most significant increased abundance following 6TG treatment, with a log_2_ foldchange of 4.70 (4.9-fold higher than the next greatest log_2_FC of all MMR proteins, RFC2), compared to MSH2 and MSH6 (0.95 and 0.67, respectively). Whilst current consensus predicates that MSH3 preferentially targets insertion/deletion loops (1-15nt) via its heteromerisation with MSH2^41^, MSH6-MSH2 heterodimer targets individual base mismatches such as that of thioguanine^42^. Our findings may suggest a more active role of MSH3 in individual base-MMR. Alternatively, 16 hours beyond the initial administration of 6TG; MSH3 may be responding to secondary replication-associated DNA-structures, rather than the initiating lesion itself.

**Fig. 3.**
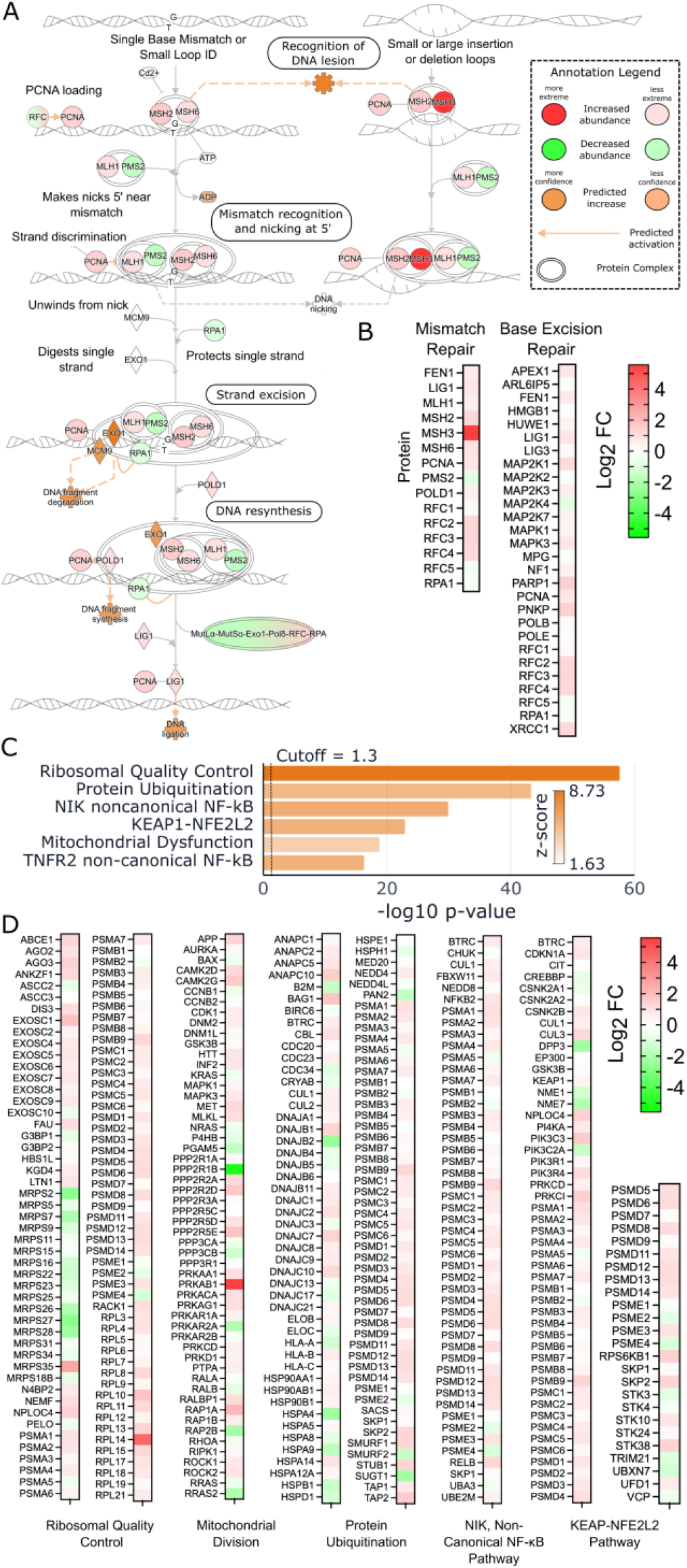
Pathway analysis from DNA-damage to secondary responses (A) Canonical DNA mismatch repair signalling pathway, overlayed with single cell proteomics protein log_2_ fold-changes (green and red proteins), with non-identified proteins represented by their predicted changes, based on global quantitative proteomics data (orange). Pathway generated in IPA (Qiagen). (B) Log_2_FC heatmap of protein-abundance changes following 6TG-treatment, for MMR and BER. (C) Bar chart representation for selected secondary pathways that were shown to be upregulated with 6TG and (D) the associated log_2_FC heatmaps for the corresponding quantified proteins.

DNA damage can promote ROS accumulation through sustained DDR signalling and disruption of mitochondrial redox homeostasis^43–45^ with oxidation of DNA-incorporated thioguanine further exacerbating genomic damage^45^. Upon 6TG induction, the primary redox homeostasis pathway: KEAP1-NFE2L2 was significantly activated; suggesting a coordinated cellular response to its increased ROS environment^45^. Complementing this finding, BER; a canonical oxidative DNA damage repair pathway^34^, was significantly enriched following 6TG-treatment, supporting a model in which 6TG induces oxidative DNA repair mechanisms, in addition to, and possibly downstream of, mismatch repair activation. Redox stress was also consistent with the significant remodelling of the mitochondrial proteome, in the context of mitochondrial stress. In particular, NADH:Ubiquinone Oxidoreductase Complex I subunits: NDUFA10, NDUFA9, NDUFB10, NDUFB3 and NDUFV1, Ubiquitinol-Cytochrome c Reductase Complex III subunits: UQCRB and UQCRC1 and Cytochrome c Oxidase Complex IV subunit: COX5A were significantly decreased (log2 FC<-1.0) in our analysis, suggesting disrupted respiratory chain function. Further ATP synthase subunits: ATP5F1B, ATP5F1D, ATP5PB and ATP5PF showed similar significantly decreased abundance profiles following 6TG-induced genotoxic stress. Corroborating this evidence of mitochondrial stress; PRKAB1, a component of the AMPK energy sensing pathway that is commonly activated during mitochondrial stress^46^, ranked as the 3^rd^ most-increased protein abundance across the entire dataset (log_2_FC: 5.05) (Fig. S3). AMPK’s role in inhibiting protein synthesis and cell growth/proliferation^47^ further supported the cellular adaptation to DNA-damage by rewiring its metabolism away from proliferative protein-synthesis signalling and towards stress responses. However, it is important to acknowledge that these hypotheses are driven by changes in protein abundances, which do not solely contribute to protein functionality. For example, AMPK pathway activity is heavily dictated by post-translational modifications and protein-protein interactions that ultimately dictate cellular behaviour^48^.

Following the archetypal progression from oxidative stress to mitochondrial and proteomic remodelling, our analysis revealed activation of multiple secondary signalling pathways indicative of inflammatory and proteostatic adaptation. Ribosomal Quality Control represented the top-ranking upregulated signalling pathway following 6TG treatment, consistent with oxidative stress–induced ribosomal stalling and disruption of translational homeostasis. Concurrent upregulation of protein ubiquitination supported a model in which stalled translational machinery and associated nascent polypeptides were targeted for proteasomal degradation to facilitate clearance of defective translation products^49^. Whilst cellular ROS-elevation from DNA-damage represents a likely stimulator of the widespread proteostatic response observed; it is important to note that 6TG also has the propensity to incorporate into RNA and can therefore influences translation in this manner. As such, this proteomic signature may reflect the combined effects of both stress-factors. Two inflammatory signalling pathways, non-canonical NF-κB inducing kinase (NIK) and TNFR2 non-canonical NF-κB^50^, were also significantly upregulated following 6TG-treatment, consistent with activation of pro-survival and inflammatory signalling programmes associated with sustained DNA damage and oxidative stress (Fig. 3B, C).

Collectively, these findings support a model in which 6TG-induced DNA-damage promotes oxidative and mitochondrial stress, leading to secondary responses including widespread proteostatic rewiring, accompanied by inflammatory signalling activation.

### Single-cell proteomics reveals changes in cell-to-cell protein-abundance heterogeneity, following DNA-damage

Previous work has established the concept of cell-to-cell heterogeneity across genetically identical cells, demonstrating substantial stochastic variability in gene expression^51^. With the advent of single-cell proteomics came the opportunity to probe the consequential impact on proteins^13,52–55^. With this in mind, we set out to investigate protein abundance heterogeneity across 3906 quantified proteins, within untreated and 6TG-treated cells, via their median absolute deviation (MAD). Across untreated U-2 OS cells, three mitochondrion-localised proteins; Isocitrate Dehydrogenase 3 Alpha (IDH3A), 3-Hydroxybutyrate Dehydrogenase Type 2 (BDH2) and Transmembrane Protein 11 (TMEM11), ranked among the ten most heterogeneous proteins based on median absolute deviation (MAD) (Fig. 4A). In contrast, the ten proteins with the most reproducible abundances, corresponding to the lowest MAD values, were all localised to the nucleus and/or cytoplasm. Indeed, comparison of the top and bottom quartiles of proteins ranked by MAD within the untreated condition, representing the most and least heterogeneous proteins in terms of cell-to-cell abundance, identified the mitochondrion as the most heterogeneous subcellular compartment. The Golgi apparatus was also selectively enriched among proteins in the top MAD quartile (Fig. S4).

**Fig. 4.**
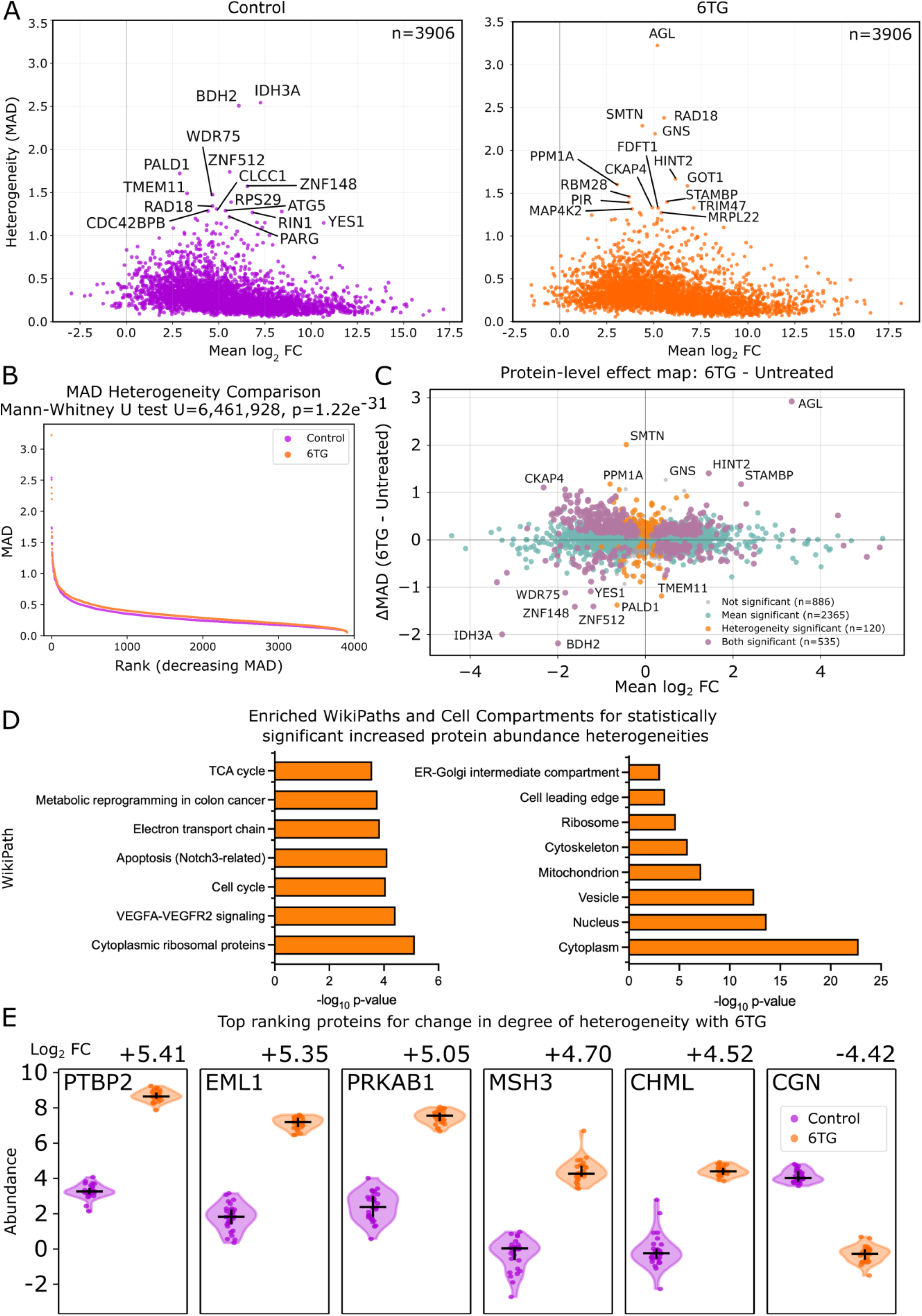
6TG-induced changes in protein abundance heterogeneity. (A) Effect map representation of cell-to-cell protein abundance heterogeneity of untreated (purple) and 6TG-treated (orange) cells, calculated as median absolute deviation (MAD), vs mean log_2_ fold-change. (B) Ranked protein heterogeneity (MAD) for all quantified proteins, for untreated (purple) and 6TG-treated (orange) cells to reveal statistically significant increase in median protein abundance heterogeneity upon 6TG treatment. (C) Effect map representation of change in protein abundance heterogeneity (measured as ΔMAD), against mean log_2_ fold-change for all quantified proteins. (D) WikiPath (left) and Gene Ontology Cellular Component (right) analysis of enriched pathways from proteins that exhibited statistically significant increased abundance heterogeneities. (E) Violin plot representation of top six ranked proteins for absolute abundance log_2_ fold-change (FC), following 6TG treatment.

We next investigated the impact of 6TG-treatment induced DNA-damage on global protein abundance heterogeneity, visualised as ranked ΔMAD for each protein in the 6TG vs control conditions (Fig. 4B, C). 6TG-treated proteins were found to show a statistically significant increase in cell-to-cell protein abundance heterogeneity. Such findings could be driven by multiple mechanistic drivers including differences in cellular 6TG uptake and DNA/RNA-incorporation^56^, and divergence of MMR-dependant biological outcomes: including ROS-adaptation, inflammation, apoptosis and degree of translational remodelling^15,57^. Further, heterogeneity may reflect differences in the temporal stage of pathway progression across individual cells. Enrichment analysis of proteins exhibiting a statistically significant change in abundance heterogeneity following 6TG-treatment (|ΔMAD| > 0.2, p_adj_ < 0.01) revealed increased heterogeneity among proteins involved in aerobic respiration (WikiPathway TCA cycle & Electron Transport Chain) and protein translation (WikiPath Cytoplasmic ribosome) (Fig. 4D). Indeed, the marked increase in heterogeneity among proteins involved in aerobic respiration was accompanied by an overall reduction in corresponding protein abundance following 6TG-induced DNA damage (Fig. S5). This pattern likely reflects secondary mitochondrial dysfunction driving suppression of oxidative phosphorylation related processes, with individual cells exhibiting variable response magnitudes and occupying different points along the stress-response trajectory. In contrast, no significantly enriched pathways were identified from proteins with reduced protein abundance heterogeneities following 6TG-treatment.

Finally, by examining the proteins exhibiting the largest 6TG-induced abundance changes through the additional lens of cell-to-cell abundance heterogeneity, we assessed whether strong population-level abundance changes were accompanied by coordinated or divergent behaviour across individual cells; 15 of the top 20 proteins (log_2_ FC) were found to undergo a reduction in abundance heterogeneity across neighbouring cells, following 6TG-treatment. In particular, PRKAB1, a mitochondrial energy metabolism regulator that is linked to mitochondrial stress^46,58^, showed a pronounced increase in abundance (log_2_ FC = 5.05, ranked 3^rd^ of 3906 proteins) alongside a significant reduction in heterogeneity (ΔMAD = −0.36; −log_10_ adjusted p-value = 2.36), consistent with the role of PRKAB1 in sensing perturbations in cellular energy balance, including altered ATP/ADP availability during mitochondrial stress^59^. Similarly, MSH3 abundance increased significantly (log_2_ FC = 4.70, ranked 4^th^ of 3906 proteins) while abundance heterogeneity was not affected by 6TG-induced DNA damage (Fig. 4E).

### Single-cell proteomics reveals bimodal response profiles for DDR and stress-response pathways following genotoxic stress

The partitioning of cellular responses into two distinct sub-populations has previously been reported, particularly in the context of DNA-damage response^60^, archetypal signalling pathways^61^ and apoptosis^62^. Thus, we next investigated evidence of distinct bimodal cellular sub-populations following genotoxic stress. To do this, we applied the Hartigan’s Dip-test^25^: a statistical approach used to assess whether a population deviates from a single continuous abundance distribution and instead exhibits evidence of multiple subpopulations. Further, we applied thresholds for sub-population fraction occupancy and valley depth, as well as ensuring the preferential fitting of abundance distribution to a 2-component Gaussian mixture model (GMM) over 1-component GMM (see *Materials and Methods*). Through this approach, several proteins were found to occupy a distinctly bimodal protein-abundance distribution that was either stimulated or abolished with 6TG-treatment (Fig. 5A, S6A).

**Fig. 5.**
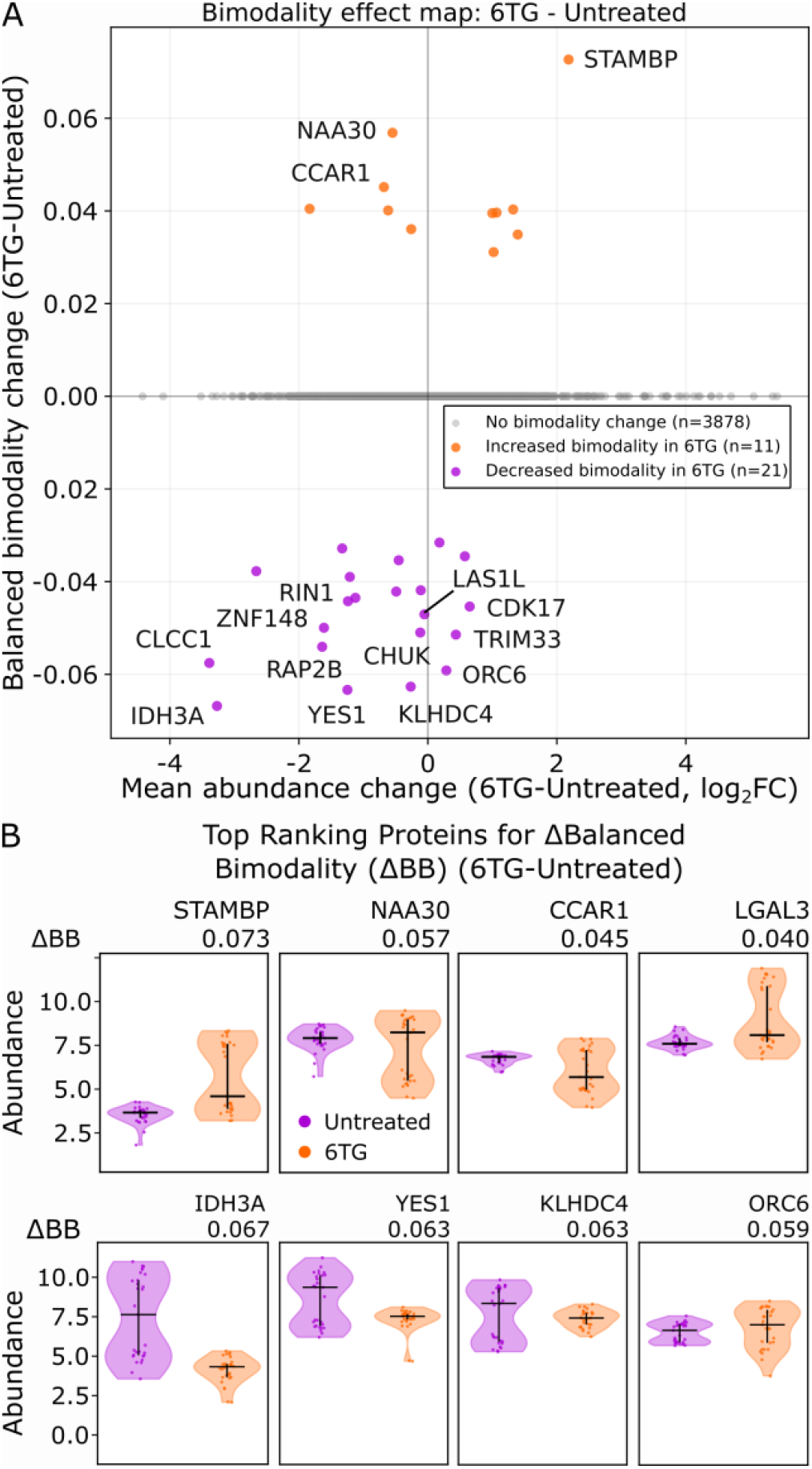
DNA-damage drives some proteins towards, and away from, bimodal functional distributions. (A) Bimodality effect map to reveal activation and abolishment of protein-abundance bimodality, upon 6TG treatment. (B) Violin plot representation of protein abundances for each cell within the untreated (purple) and 6TG-treated (orange) cellular population, including the top 4 ranked proteins for activation and suppression of bimodality.

STAMBP and LGALS3, a deubiquitinating enzyme linked to regulation of protein-trafficking/lysosomal degradation^63^ and an upstream activator of NF-κB signalling^64,65^, respectively, exhibited the bifurcation of abundance responses following 6TG-induced DNA damage. One cellular subpopulation showed little to no change in protein abundance, whereas a second subpopulation displayed a pronounced abundance increase (Fig. 5B). Also displaying this behaviour was PMS1, a canonical mismatch repair protein that has been implicated in promoting somatic CAG-repeat expansion, a process strongly associated with modification of age at onset in Huntington’s disease^66^ (Fig. S6B). Conversely, N-acetyltransferase C subunit NAA30, whose depletion is associated with mitochondrial damage^67^, exhibited the opposite pattern, with one cellular subpopulation showing a significant reduction in abundance while the remaining cells displayed little or no change (Fig. 5B). Taken together, these findings may suggest the existence of a cellular stimulatory threshold beyond which synthesis of primary DNA damage response related proteins and secondary oxidative stress-related proteins may be regulated, either positively (PMS1, STAMBP, LGALS3) or negatively (NAA30, CCAR1).

Intriguingly, several proteins exhibited a bimodal abundance distribution across the untreated U-2 OS cells, with this intrinsic bifurcation abolished upon 6TG-treatment. For example, IDH3A, the mitochondrial α- subunit of NAD^+^-dependent isocitrate dehydrogenase in the tricarboxylic acid cycle, and YES1, an Src-family tyrosine kinase associated with proliferative signalling, both lost their bimodal abundance distributions following 6TG treatment, converging toward the lower-abundance subpopulation. Indeed, this may reflect suppression of mitochondrial oxidative phosphorylation and cell division, during DNA-damage related stress response, although further experiments are required to validate this. Interestingly, IKKα (CHUK), a key component for the activation of non-canonical NF-κB signalling, also exhibited an abolishment of bimodality, with cells converging towards the higher abundance state upon 6TG-treatment (Fig. S6B). The activation of NIK and TNFR2 pathways (Fig. 3C) support this homogenisation towards non-canonical NF-κB inflammatory signalling, as a secondary response to DNA-damage. Taken together, the collapse of a pre-existing protein-abundance bimodality towards a more homogeneous distribution after DNA damage may reflect the stress induced convergence of some proteins, relating to primary- and secondary-responses, towards a more distinct cellular state.

Collectively, these findings suggest that, while 6TG-induced DNA damage remodels mean protein abundance, the enhanced resolution afforded by SCP reveals distinct cell-to-cell responses that are obscured at the population level. The emergence of high- and low-abundance subpopulations for some proteins, alongside convergence towards defined abundance states for others, highlights substantial heterogeneity in the cellular response to genotoxic stress. Such behaviour may reflect both threshold-dependent activation of stress-response programmes and coordinated convergence towards common DNA damage, inflammatory and metabolic states. Importantly, further work to independently validated these findings, using targeted and orthogonal biochemical approaches, is necessary to confirm the biological significance and mechanistic basis of the observed bimodal protein-abundance distributions. Such studies will also be necessary to determine the extent to which additional sources of cellular heterogeneity, including variation within assigned cell-cycle states, contribute to the observed bimodal behaviour.

## CONCLUSIONS

In this study, we applied single-cell proteomics to characterise proteome remodelling following 6-thioguanine–induced DNA damage response in U-2 OS cells, whilst specifically controlling for confounding heterogeneity arising from dead-cell contamination, suboptimal proteomics sampling, and cell-cycle variation. By focusing our analysis on a Seurat inferred G1-phase cellular population undergoing replication-associated genotoxic stress, we resolved coordinated activation of canonical mismatch repair and base excision repair pathways alongside extensive secondary proteome remodelling associated with oxidative stress, mitochondrial dysfunction, inflammatory signalling, and proteostasis. Within the canonical DNA Mismatch Repair pathway; MSH3 represented the 4^th^ greatest increase in abundance across all quantified proteins, underscoring its significance to the mismatch repair machinery. Reinforcing the secondary response biology that is propagated by DNA-damage; PRKAB1, a regulator of mitochondrial energy metabolism that is linked to mitochondrial stress^58^, exhibited the 3^rd^ largest increase in abundance following 6TG-treatment. Together, these findings support a model in which 6-thioguanine-induced DNA lesions propagate beyond primary DNA repair signalling to drive broader stress-adaptive cellular programmes linked to ROS-accumulation (as demonstrated by the significantly increased KEAP1-NFE2L2 oxidative stress response pathway)^5,68^.

Importantly, single-cell resolution enabled interrogation not only of average protein abundance changes, but also of the distribution of protein-abundances across neighbouring cells. Quantitative heterogeneity analysis revealed widespread increases in protein abundance variability following genotoxic stress, particularly across oxidative phosphorylation and the electron transport train within the mitochondrion. Against this backdrop of increased abundance heterogeneity; PRKAB1’s strikingly increased abundance was accompanied by a significant reduction in cell-to-cell abundance heterogeneity; demonstrating the robustly reproducible response of this mitochondrial energy metabolism sensor upon 6TG-mediated DNA damage.

Beyond the normal distribution of protein abundances around a mean value across individual cells, our workflow revealed both the emergence and abolishment of high and low protein abundances following 6TG-treatment. Exemplifying this, canonical MMR response protein PMS1, and proteostasis-& inflammation-related proteins STAMBP and LGALS3, respectively, exhibited the establishment of bifurcation of abundances to high and low populations upon DNA-damage induced 6TG-treatment. Conversely, abolishment of this intrinsic bimodality for several proteins upon 6TG-treatment was also observed. Collectively, these findings support a model in which 6TG-induced DNA damage drives the bifurcation of some proteins, reflective of differential engagement of pathways upon DNA-damage, whilst some proteins undergo the homogenisation of their abundance from an intrinsic bimodal distribution in response to this genotoxic stress. Understanding the origins of these heterogeneous and bimodal responses will be essential for resolving the divergent and convergent proteome trajectories that emerge during prolonged genotoxic stress.

This work demonstrates the utility of SCP for resolving mechanistic features of DNA damage response that remain obscured in bulk-cell measurements. By integrating proteome-wide abundance profiling with quantitative analysis of cellular heterogeneity and bimodality, our study reveals that, beyond altering protein abundances, genotoxic stress reshapes the distribution of cellular proteomic states across genetically identical individual cells, to varying degrees. These findings establish single-cell proteomics as a powerful framework for dissecting subpopulation-specific stress responses to provide a broader foundation for investigating heterogeneous cellular responses, associated with genome instability. Further investigation of the mechanistic basis of cell-to-cell protein abundance heterogeneity and bimodality, together with their functional consequences, will reveal how these distinct proteomic states influence adaptation to genotoxic stress and ultimately shape cell fate.

## Supporting information

Differential pathways for pre- vs-post filtering

Protein abundances post filtering

Heterogeneity analysis

Bimodality analysis

Protein abundances pre filtering

## ASSOCIATED CONTENT

### Supporting Information

*The Supporting Information is available free of charge on the ACS Publications website*.

*Matrix of protein abundances, pre-filtering (csv)*

*Matrix of protein abundances, post-filtering (csv)*

*Signalling pathway analysis for pre- and post-filtered proteomics data (csv)*

*Heterogeneity data corresponding to heterogeneity parameters for each protein (csv)*

*Bimodality data corresponding to bimodality parameters for each protein (csv)*

*GitHub code accessible via: https://github.com/kish-adoni/SCP_heterogeneity-and-bimodality or via* https://github.com/orgs/ThalassinosLab/repositories

*Proteomics RAW data is available via Proteome Xchange Proteome Xchange PRIDE identifier: PXD074838*

*Reviewer Username:*

*Reviewer Password: PbgfHTJS7XZF*

## AUTHOR INFORMATION

## Author Contributions

Conceptualisation: K. R. A., G. H. C., R. Z. C., K. T.

Methodology: G. H. C., R. Z. C., J. H.

Software: K. R. A., J. E. D., D. T. C.

Validation: K. R. A., G. H. C., J. E. D., D. T. C., J. H., R. Z. C

Formal Analysis: K. R. A, G. H. C., J. E. D., D. T. C.

Investigation: G. H. C., J. H.

Data Curation: K. R. A., J. E. D.

Visualisation: K. R. A, J. E. D.

Writing – Original Draft: K. R. A., G. H. C., J. E. D.

Writing-Reviewing and Editing: K. R. A., G. H. C., J. E. D., D. T. C., J. H., J. K., K. T.

Project Administration: K. T. Funding Acquisition: K. T.

### Funding Sources

Wellcome Trust Multiuser Equipment grant 221521/Z/20/Z (K.T). Wellcome Trust Collaborative Award in Science 209250/Z/17/Z (K.T). CHDI Foundation (K. R. A, G. H. C)

## ABBREVIATIONS

MMR: Mismatch repair
DDR: DNA damage response
BER: Base excision repair
SCP: Single cell proteomics
6TG: 6-thioguanine
ROS: Reactive oxygen species
MAD: Median absolute deviation
FC: Fold-change
TCA: Tricarboxylic acid.

