## Supplementary Figures for "Single-Cell Proteomics Reveals Heterogeneous and Bimodal Proteome Responses to DNA Damage"

**Supplementary Information: Single-Cell Proteomics Reveals Heterogeneous and Bimodal Proteomic Responses to DNA Damage**

Kish R. Adoni^†1^, Georgina H. Charlton^†1^, Jonathan E. Ditcham^1^, Declan Cook^1^, Jenny Ho^3^, Joanna Kirkpatrick^3^, Riccardo Zenezini Chiozzi^1,4^, Konstantinos Thalassinos^1,2,4^*

1 Research Department of Structural and Molecular Biology, Division of Biosciences, Gower Street, London WC1E 6BT, United Kingdom

2 School of Natural Sciences, Birkbeck College, London, WC1E 7HX, United Kingdom

3 Thermo Fisher Scientific, Hemel Hempstead, UK.

4 University College London Mass Spectrometry Science Technology Platform, Division of Biosciences, University College London, London, UK

†*These authors contributed equally to this work*

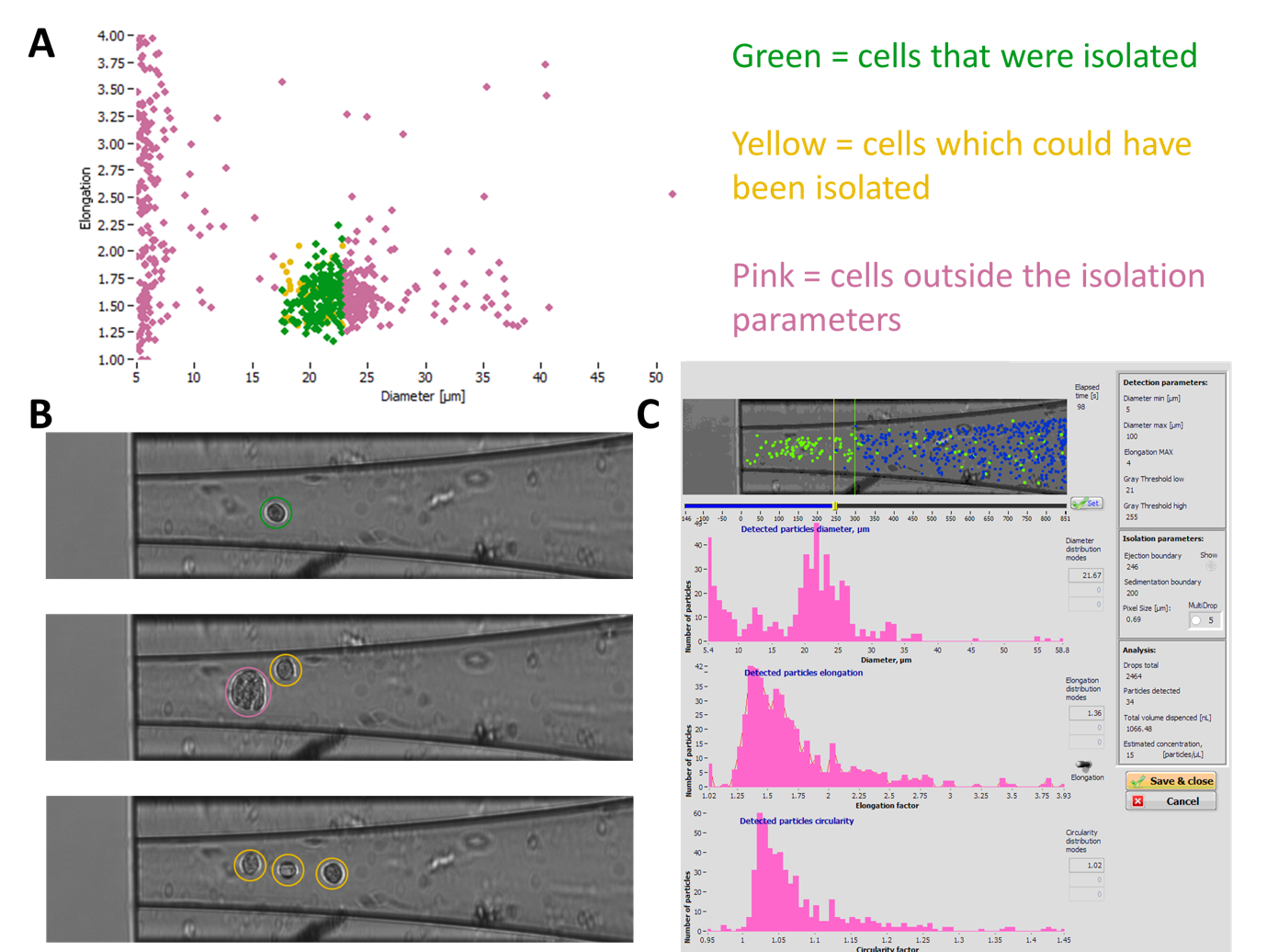

Fig. S1: Single-cell isolation. (A) Plot to demonstrate cell selection criteria, with green dots selected for onward sample preparation and MS-analysis. (B) CellenONE microscope images of cells travelling through Piezo Dispense Capillary (PDC). Green annotated cell (top) represents a viable cell for analysis, whereas pink annotated cell (middle) represents a morphological parameters that were outside pre-defined thresholds and yellow annotated cells (bottom) represents cells that were within parameter thresholds, but were not isolated as they would be isolated with other cells. (C) Screenshot from CellenONE instrument computer, with selected detection and isolation parameters (right) and histograms for number of particles at specific ranges of diameter, elongation factor and circularity factor.

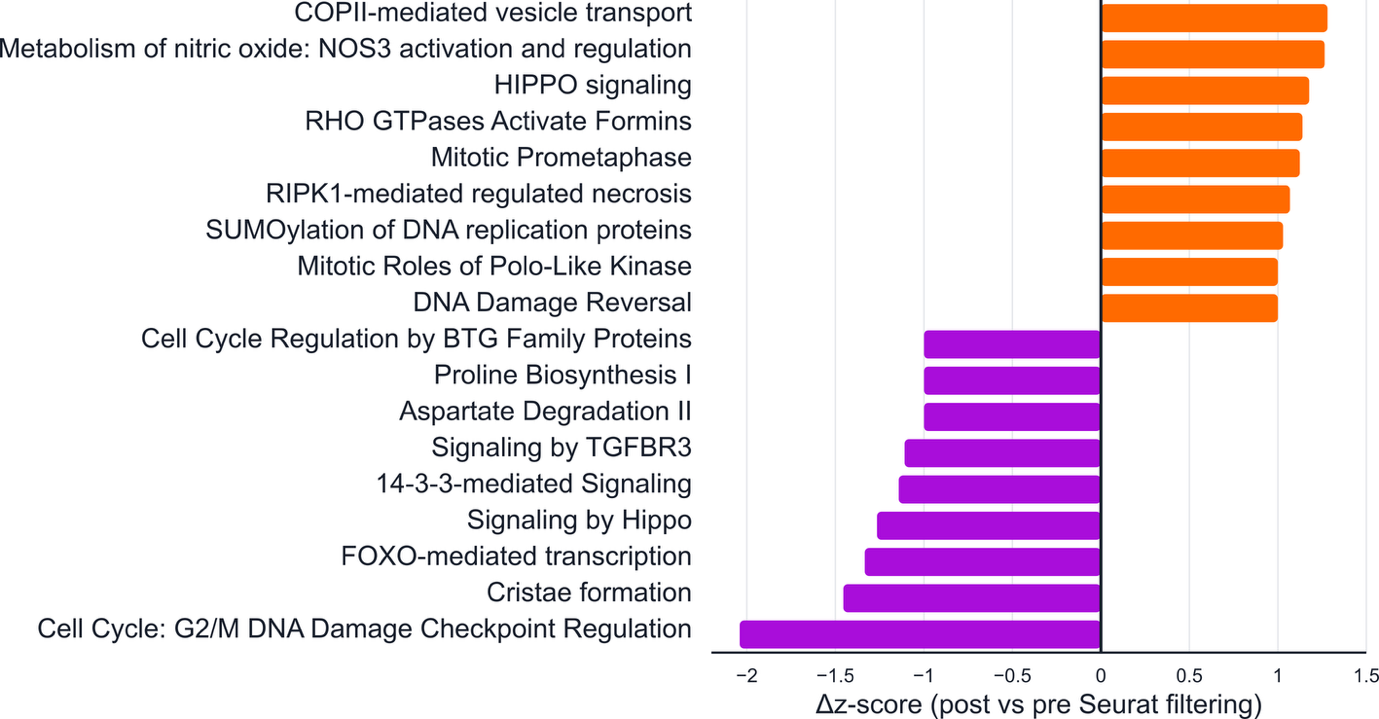

Fig. S2: Difference in z-score of pathways following Seurat cell-cycle filtering, for top pathways based on absolute z-score.

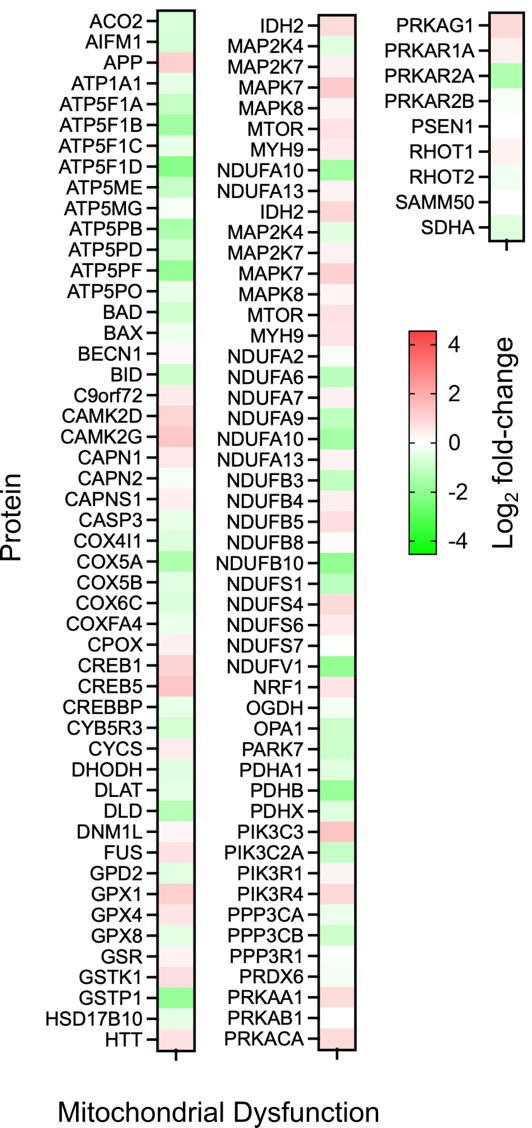

Fig. S3: Protein abundance heatmap for all proteins relating to mitochondrial dysfunction; following 6TG treatment.

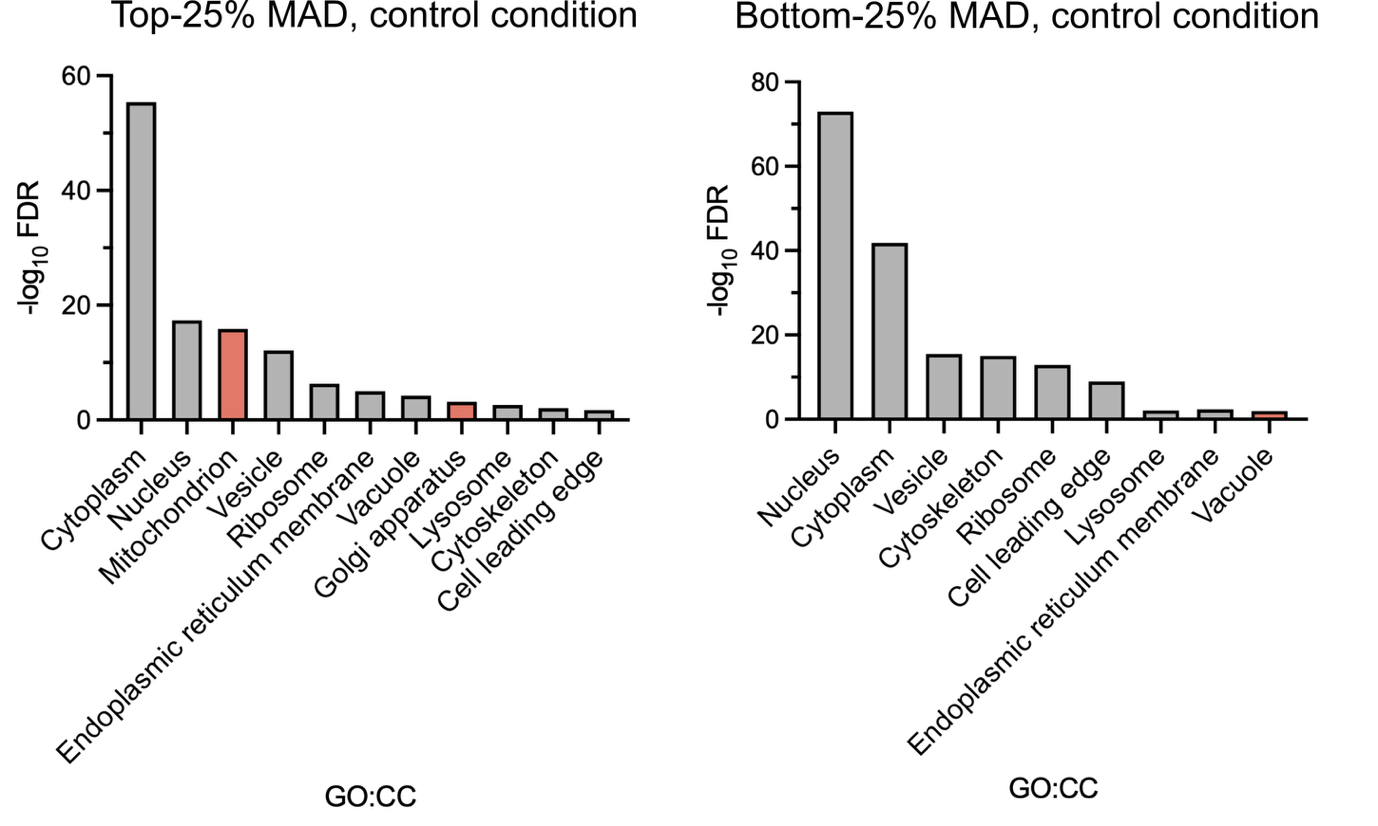

Fig. S4: Gene Ontology: Cell Compartment (GO:CC) enriched compartments for top-quartile most heterogenous proteins (left) and bottom quartile least heterogeneous proteins (right), based in median absolute deviation (MAD) in untreated control condition of U-2 OS cells. Red bars indicate unique presence of that compartment in top- or bottom-quartile.

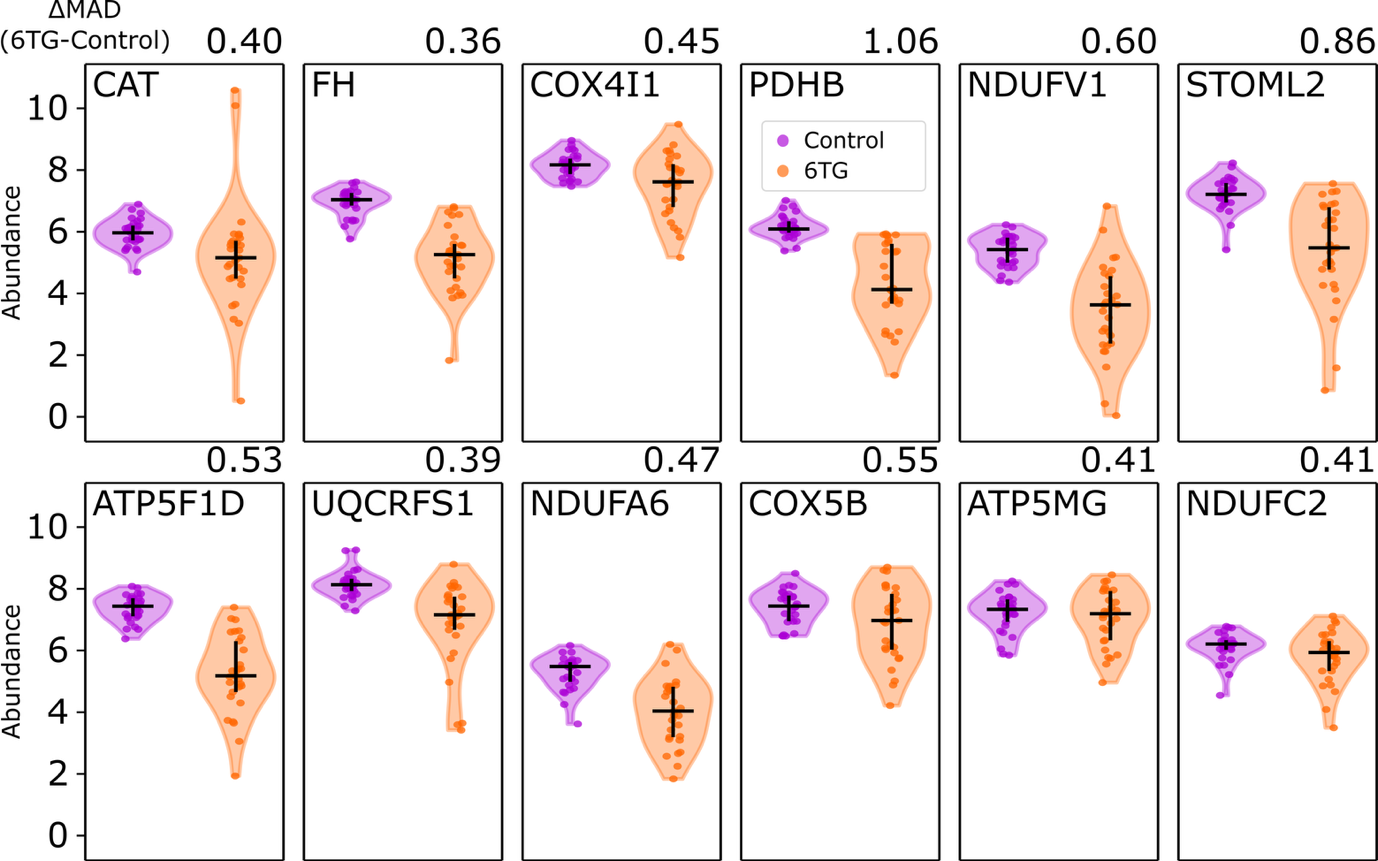

Fig. S5: Protein abundance violin plots for aerobic respiration related proteins, demonstrating their increased heterogeneity, concomitant with reduced abundance upon 6TG-treatment induced DNA-damage.

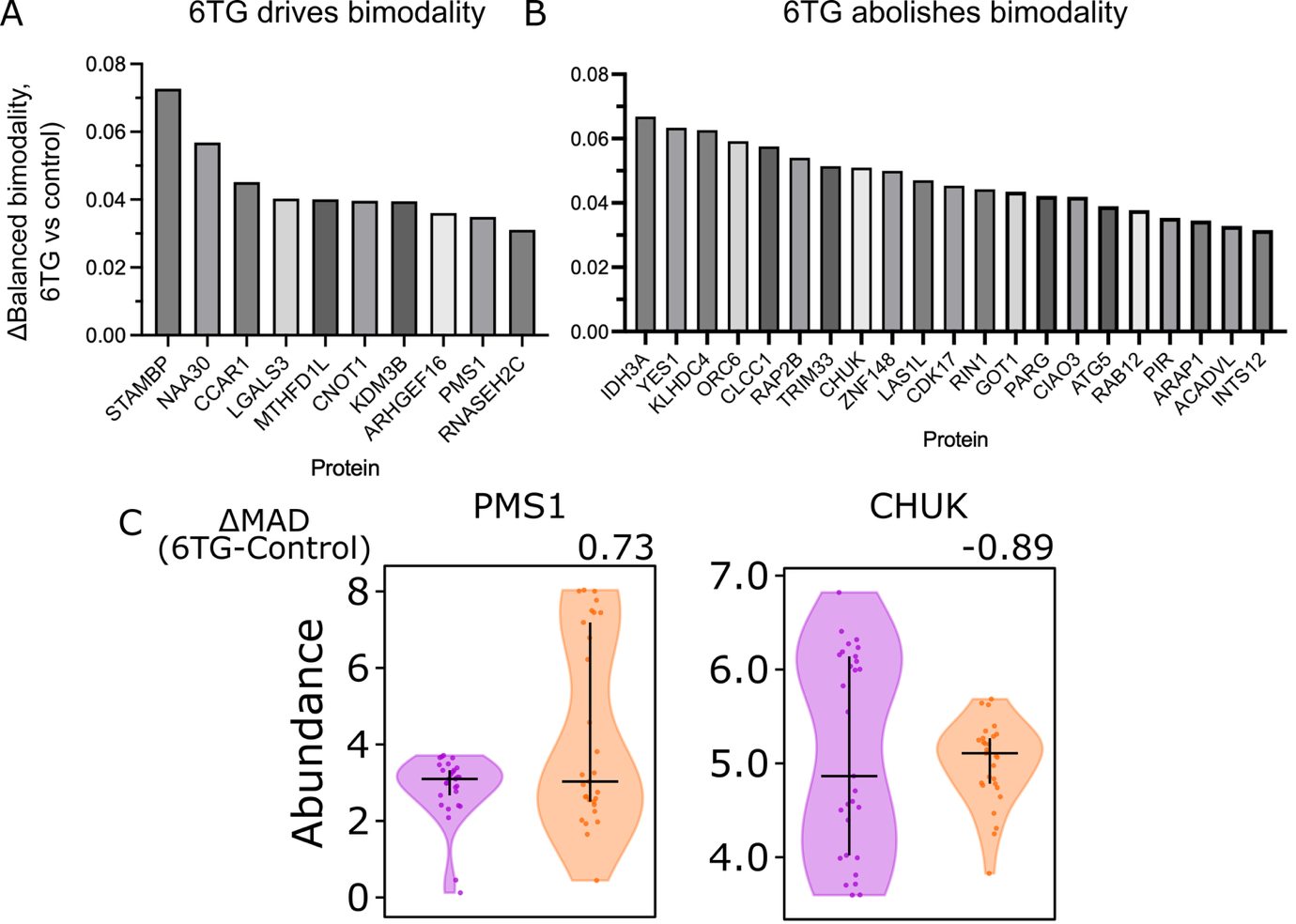

Fig. S6: Bimodality analysis. Proteins with (A) acquisition of bimodal abundance behaviour and (B) abolishment of bimodal behaviour, upon 6TG-treatment. (C) Protein abundance violin plots to demonstrate gain of bimodality for PMS1 and abolishment of bimodality for CHUK (IKKα).
